# Increased positive selection in highly recombining genes does not necessarily reflect an evolutionary advantage of recombination

**DOI:** 10.1101/2024.01.16.575829

**Authors:** Julien Joseph

## Abstract

It is commonly thought that the long-term advantage of meiotic recombination is to dissipate genetic linkage, allowing natural selection to act independently on different loci. It is thus theoretically expected that genes with higher recombination rates evolve under more effective selection. On the other hand, recombination is often associated with GC-biased gene conversion (gBGC), which theoretically interferes with selection by promoting the fixation of deleterious GC alleles. To test these predictions, several studies assessed whether selection was more effective in highly recombining genes (due to dissipation of genetic linkage) or less effective (due to gBGC), assuming a fixed distribution of fitness effects (DFE) for all genes. In this study, I directly derive the DFE from a gene’s evolutionary history (shaped by mutation, selection, drift and gBGC) under empirical fitness landscapes. I show that genes that have experienced high levels of gBGC are less fit and thus have more opportunities for beneficial mutations. Only a small decrease in the genome-wide intensity of gBGC leads to the fixation of these beneficial mutations, particularly in highly recombining genes. This results in increased positive selection in highly recombining genes that is not caused by more effective selection. Additionally, I show that the death of a recombination hotspot can lead to a higher *dN/dS* than its birth, but with substitution patterns biased towards AT, and only at selected positions. This shows that controlling for a substitution bias towards GC is therefore not sufficient to rule out the contribution of gBGC to signatures of accelerated evolution. Finally, although gBGC does not affect the fixation probability of GC-conservative mutations, I show that by altering the DFE, gBGC can also significantly affect non-synonymous GC-conservative substitution patterns.

## Introduction

Meiotic recombination is a key cellular process with major consequences for evolution. In the vast majority of sexually reproducing species, the formation of a crossing over is required for the proper segregation of homologs during meiosis (Baker et al., 1976; Davis and Smith, 2001; Pardo-Manuel de Villena and Sapienza, 2001; Gerton and Hawley, 2005). Failure of this process often leads to meiotic arrest or the formation of aneuploid gametes with missing or extra chromosomes (Hassold et al., 2007; Handel and Schimenti, 2010; Székvölgyi and Nicolas, 2010; Brick et al., 2012; Mihola et al., 2019).

From an evolutionary perspective, the most commonly cited hypothesis for the long-term maintenance of recombination is its effect on genetic linkage (Felsenstein, 1974; Otto and Barton, 1997; Otto and Lenormand, 2002; Keightley and Otto, 2006). By creating new combinations of alleles, recombination events dissociate the selective pressures exerted on different loci, dissipating the so-called Hill-Robertson interference (Hill and Robertson, 1966; Smith and Haigh, 1973; Charlesworth et al., 1993; Roze and Barton, 2006). A large number of theoretical studies have explored conditions under which recombination can have a positive, negative or no effect on the efficacy of natural selection. As a result, it is now generally accepted that in populations of finite size and where new mutations are mostly deleterious, increased recombination rates enhance the efficacy of both purifying and positive selection (Felsenstein, 1974; Hickey and Golding, 2018; Roze, 2021). To test this theoretical result, several empirical studies tried to quantify signatures of selection across regions of different recombination rate, with mixed results (Bullaughey et al., 2008; Gossmann et al., 2014; Hussin et al., 2015; Bolívar et al., 2016; Castellano et al., 2016; Corcoran et al., 2017; Rousselle et al., 2019; Murga-Moreno et al., 2019; Grandaubert et al., 2019; Castellano et al., 2020; Cavassim et al., 2021). Some studies claim a beneficial effect of recombination (Gossmann et al., 2014; Hussin et al., 2015; Castellano et al., 2016; Corcoran et al., 2017; Murga-Moreno et al., 2019; Castellano et al., 2020), some claim no effect of recombination on the efficacy of selection (Bullaughey et al., 2008), and others claim a deleterious effect (Necşulea et al., 2011; Lachance and Tishkoff, 2014; Bolívar et al., 2016).

One of the reasons for these mixed results is that recombination has another effect on genomes that has been largely overlooked in theoretical studies: GC-biased gene conversion (gBGC) (Brown and Jiricny, 1987; Duret and Galtier, 2009). The very mechanism of recombination requires hybridisation between the two single-stranded parental DNAs. If an individual is heterozygous at this position, this will cause a mismatch in the resulting double-stranded DNA (heteroduplex) that can be repaired in the direction of either allele. One allele thus converts the other and this phenomenon is therefore called gene conversion (Winkler, 1930) (reviewed in Roman (1985)). In many Eukaryotes including vertebrates, plants and fungi, recombination-associated gene conversion is biased towards GC alleles (Pessia et al., 2012; Galtier et al., 2018). The evolutionary consequences of this mechanism is the rapid spread of GC alleles in regions of high recombination rates, eventually leading to their rapid fixation in the population.

As the main methods of inference of positive selection rely on the detection of rapid fixation events (McDonald and Kreitman, 1991), the pervasive fixation of GC alleles acts as a confounding factor (Galtier et al., 2009; Ratnakumar et al., 2010). Ratnakumar et al. (2010) estimated that *∼*20% of protein-coding genes showing a rapidly increasing *dN/dS* ratio in the human branch were likely to be the result of the birth of a gBGC hotspot, and did not correspond to adaptation to the new human environment as previously hypothesized (Kosiol et al., 2008). This proportion was estimated using the fact that increased gBGC skews substitution patterns towards GC, even in non coding sequence, which is not expected under positive selection only (Ratnakumar et al., 2010).

However, it is not clear whether contrasting positive selection across genes with different recombination rates is a suitable way to test for the beneficial effect of recombination, even when controlling for the rapid fixation of GC alleles. Indeed, even true positive selection in highly recombining genes is not necessarily the result of a higher selection efficacy. In fact, the rate of positive selection is the product of the number of beneficial mutations per generation, times their fixation probability (Kimura, 1962). By interpreting increased positive selection as a sign of increased selection efficacy, a strong hypothesis is implicitly made: genes that evolved under different recombination rates have the same number of opportunities for beneficial mutations. Alternatively stated, the above-mentioned empirical studies implicitly (or sometimes explicitly) assume an invariant distribution of fitness effects (DFE) of new mutations across recombination rate categories (Bullaughey et al., 2008; Gossmann et al., 2014; Hussin et al., 2015; Bolívar et al., 2016; Castellano et al., 2016; Corcoran et al., 2017; Rousselle et al., 2019; Grandaubert et al., 2019; Murga-Moreno et al., 2019; Castellano et al., 2020; Cavassim et al., 2021).

The DFE of new mutations is a key parameter in evolution, and largely depends on the fitness landscape. Indeed, in a gene that is strongly selectively constrained, most new mutations will be deleterious. Conversely in a non-functional pseudogene, all mutations will be neutral. Moreover, the DFE will be affected by the position of a sequence in its fitness landscape. Indeed, in a gene that is close to its fitness optimum, most mutations will be deleterious. Conversely, in a gene that is far from its fitness optimum, many mutations will be beneficial. In a more formal phrasing, the DFE can be seen as the local derivative of the fitness landscape at the position of a sequence in it, and can be computed as such (Martin and Lenormand, 2006).

As the fitness landscape is usually difficult to access, it is a common practice in population genetics to consider the DFE as a fixed parameter (e.g. Bolívar et al. (2016) and Corcoran et al. (2017) for theoretical models of gBGC), even though the DFE is supposed to depend on the other parameters of the model. In particular, the position of the sequence in a fitness landscape is driven by natural selection, mutation, drift and gBGC.

In this paper, I used a simple mutation-selection model to compute the expected DFE in genes that have evolved under different recombination rates, under a collection of experimental and empirical fitness landscapes. I show that by promoting the fixation of many deleterious mutations, gBGC drives sequences away from their optimal fitness, which directly creates more opportunities for beneficial back-mutations and skews the DFE towards positive values in highly recombining genes. Upon a decrease of the repair bias towards GC, or a decrease of the genome-wide recombination rate (keeping the recombination landscape constant), those beneficial back-mutations eventually get fixed by positive selection. This creates a true signal of positive selection in highly recombining genes that does not imply either higher selection efficacy or increased adaptation. Finally, I show that the death of a long-lived recombination hotspot can generate an even higher *dN/dS* than its birth. Of note, in this case, substitution patterns are biased towards AT, and only at selected positions. Importantly, this increase in *dN/dS* is due to positive selection, but is not a signature of an adaptation to changing environments. Therefore, these genes potentially add to the list of false positives when using accelerated evolution or signatures of positive selection to test for adaptive evolution (Hartl and Taubes, 1996; Galtier and Duret, 2007; Mustonen and Lässig, 2009; Jones et al., 2017).

## Results

### Impact of gBGC on the equilibrium DFE of new mutations

As stated in the introduction, the DFE of new mutations depends both on the fitness landscape, and on the position of a given individual in it. A total of six amino-acid fitness landscapes were used in this study: one mammalian fitness landscape obtained by fitting a mutation-selection model to multi-species alignments of 14,509 protein-coding genes in Latrille et al. (2023), four fitness landscapes obtained from deep mutational scanning (DMS) experiments in Influenza (Thyagarajan and Bloom, 2014; Doud et al., 2015), *E*.*coli* (Stiffler et al., 2015), and *S*.*cervisae* (Kitzman et al., 2015), and a concatenate of the four previous DMS fitness landscapes. As the fitness landscapes are fixed, I therefore study selection dynamics in the absence of adaptation (Hartl and Taubes, 1996; Mustonen and Lässig, 2009; Jones et al., 2017). Moreover, because the fitness landscapes are site-specific, I thus neglect epistatic interactions. The consequences of this latter assumption are discussed below. The position of a sequence in these fitness landscapes depends on its evolutionary history. In particular, this position will result from an equilibrium between mutation, selection, drift and gBGC (Nagylaki, 1983; Hartl and Taubes, 1998; Sella and Hirsh, 2005; Mustonen and Lässig, 2009). Using a mutation-selection model, one can compute the equilibrium frequency for each codon at each site for all the fitness landscapes (details in methods).

At mutation-selection-drift-gBGC equilibrium, by sampling all possible mutations, weighted by the equilibrium frequency of the ancestral codon and their mutation rate, and associating them with their fitness effect, one directly obtains a DFE of new mutations (see details in methods). I computed the DFE separately for WS (from AT ↦ GC) and SW (from GC ↦ AT) mutations, using a population-scaled gBGC coefficient (*B* = 4*N*_*e*_*b*) of 0 (no gBGC) or 2. For all the DFEs presented in this study, I considered only non-synonymous mutations. To be able to compare the results obtained on different fitness landscapes, I rescaled them such that the mean population-scaled selection coefficient from the fittest codon (*S* = 4*N*_*e*_*s*) is 20. This ensures that the selective pressure on protein is high enough such that most sites are little influenced by gBGC, as expected in reality (Duret and Galtier, 2009).

First, I computed equilibrium frequencies with a Jukes Cantor mutation matrix (which assumes that all bases have equal mutation rates). It appears that the positive part of the DFE is almost identical between WS and SW mutations (Fig.1 and Table 1). However, even in the absence of gBGC or mutational bias, there appears to be slightly more opportunity for deleterious mutations towards GC than towards AT (Fig.1). This can be partly explained by the structure of the genetic code: a majority of amino-acids have an obligatory A or T in first or second position of codons. Indeed, when randomly switching amino-acids in the fitness landscape of a given site, there are still more deleterious mutations towards GC (Fig. S1).

**Table 1:** Proportion of beneficial mutations under *B* = 0 and a Jukes Cantor (JC) mutation matrix, *B* = 0 and a human mutation matrix (HM) and *B* = 2 and a human mutation matrix.

|  | B = 0 JC |  |  | B = 0 HM |  |  | B = 2 HM |  |  |
| --- | --- | --- | --- | --- | --- | --- | --- | --- | --- |
| Fitness landscape | All | WS | SW | All | WS | SW | All | WS | SW |
| Mammalian | 1.33% | 1.13% | 1.48% | 1.37% | 1.62% | 1.11% | 1.64% | 0.35% | 2.97% |
| Concatenated DMS | 3.04% | 2.68% | 3.33% | 3.02% | 3.66% | 2.54% | 3.22% | 0.79% | 5.39% |
| NP (Influenza) | 0.61% | 0.68% | 0.54% | 0.72% | 1.30% | 0.42% | 0.71% | 0.15% | 1.27% |
| HA (Influenza) | 1.42% | 1.15% | 1.74% | 1.51% | 1.7% | 1.38% | 1.73% | 0.29% | 3.35% |
| Gal4 (*S.cervisiae*) | 1.57% | 1.33% | 1.83% | 1.37% | 1.81% | 0.97% | 1.93% | 0.43% | 3.84% |
| β-lactamase (*E.coli*) | 11.9% | 10.8% | 12.4% | 11.6% | 12.7% | 10.3% | 12.11% | 4.47% | 16.2% |

**Figure 1.**
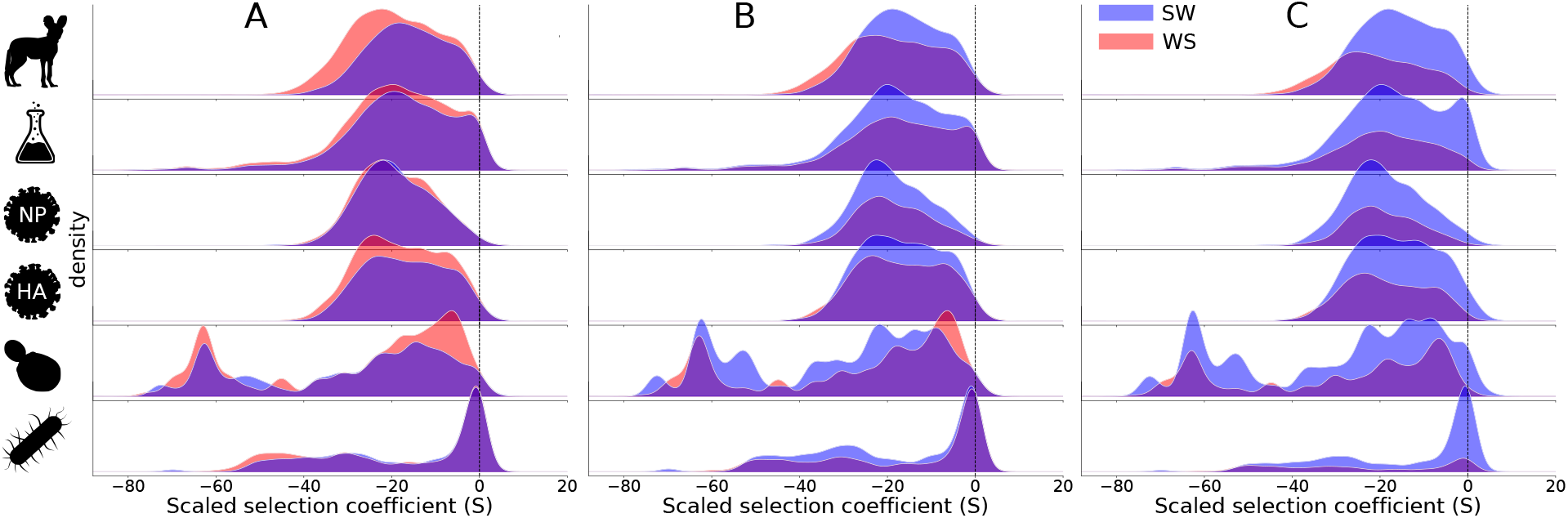
Distribution of fitness effects of new mutations at equilibrium separately for WS (red) and SW (blue) mutations. Equilibrium frequencies were computed for each fitness landscape under A) *B* = 0 and a JC mutation matrix, B) *B* = 0 and a human mutation matrix and C), *B* = 2 and a human mutation matrix. From top to bottom: 2000 sites randomly sampled from the mammalian fitness landscapes, the concatenate of the DMS fitness landscapes (1389 sites), the fitness landscape of the influenza protein NP (498 sites), the influenza protein HA (564 sites), the *S*.*cervisae* protein Gal4 (64 sites) and the *E*.*coli* protein *β*-lactamase (263 sites).

Using a realistic human mutation matrix, where mutations are typically biased towards AT, the number of SW mutations increases as expected (Fig.1). Importantly, this increase is not uniform along the DFE, and mostly concerns the deleterious compartment (Fig.1). Indeed, beneficial mutation towards AT have already been fixed, and only mutations that are so deleterious that they cannot be fixed remain as mutational opportunities. Although the number of expected SW and WS beneficial back mutations is roughly the same, it is important to note that the proportion of beneficial WS mutations among all WS is higher than the proportion of beneficial SW mutations among all SW (Table 1). This means that WS mutations are on average more beneficial than SW ones. This result has already been found with a similar approach by Latrille and Lartillot (2022) and Kaj et al. (2023).

When the sequence has evolved under gBGC (*B* = 2, corresponding to a region of high recombination rate in the human genome estimated by Glémin et al. (2015)), the SW mutation rate is still higher because of the mutational bias, but given that gBGC has increased GC content, the number of SW mutational opportunities is also higher. This results in even more SW mutations (Fig.1). Moreover, the DFE of WS mutations is skewed towards negative values, while the DFE of SW mutations is skewed towards positive values (Fig.1 and Table 1). Indeed, at equilibrium, GC alleles that were slightly deleterious have been fixed because of gBGC, and SW mutations are therefore more beneficial on average (Table 1). Of note even if SW mutations are more beneficial, they do not fix more often because of gBGC, and remain as mutational opportunities. Conversely, most WS mutations that were slightly advantageous have already been fixed by both positive selection and gBGC. Therefore, only WS mutations that cannot fix because they are deleterious enough to be wiped out by purifying selection even when helped by gBGC remain available as mutational opportunities (Table 1). It is also important to note that when *B* = 2, one always expects more beneficial back-mutations in total than without gBGC (Table 1). Interestingly, Glémin (2010) showed that gBGC cancels the fixation load induced by the mutational bias towards AT (*λ*) when *B* = *ln*(*λ*), and that this value does not depend on the distribution of selective effects (which is analogous to a fitness landscape in bi-allelic models). For the human mutational bias, it means a *B* close to 0.7. However, this is assuming that half optimal states are GC and half are AT. As shown here, the genetic code (and potentially other factors) increases the proportion of AT optimal states, which already alleviates the load induced by the mutational bias. In this case, the value of *B* that perfectly compensate the mutational bias is expected to be even lower than 0.7.

### Impact of gBGC fluctuations on selection dynamics

Similarly to the DFE of new mutations, by sampling all possible mutations weighted by the equilibrium frequency of the ancestral codon and their substitution rate, and by associating them to their fitness effect, one directly obtains a DFE of substitutions (see details in methods). I computed this DFE of substitutions at equilibrium for *B* = 0 and *B* = 2 using the realistic human mutation matrix. As the results are almost identical for all fitness landscapes, I only represented results obtained on the concatenate of DMS fitness landscapes in Fig.2, and presented the other ones in Supplementary Fig. S2. For *B* = 0, there are as many beneficial substitutions as deleterious ones (Fig.2A and Fig. S2). Additionally, there are as many WS substitutions as SW ones (Fig.2A and Fig. S2). This is expected since the sequence is at equilibrium both for fitness and base composition. For *B* = 2, the exact same conditions are met: there are as many beneficial substitutions as deleterious ones and there are as many WS substitutions as SW ones, again, the sequence is at equilibrium (Fig.2B and Fig. S2). However, as hinted from the DFE of new mutations, WS substitutions are more often deleterious, while SW substitutions are more often beneficial (Fig.2B and Fig. S2). To gain insight on selection dynamics out of equilibrium, I computed the equilibrium frequencies of codons under *B* = 2, and computed the substitution rates from this equilibrium sequence either with *B* = 1 or *B* = 3 (Fig.2C&D and Fig. S2). When gBGC increases (*B* = 2 *1↦ B* = 3), there is an increase in the fixation of deleterious GC alleles (Fig.2C and Fig. S2), as predicted by theoretical models, and observed empirically in diverse organisms (Nagylaki, 1983; Bengtsson, 1990; Galtier et al., 2009; Glémin, 2010; Necşulea et al., 2011; Bolívar et al., 2016; Rousselle et al., 2019). Conversely, when gBGC decreases (*B* = 2 ↦ *B* = 1), there is an equivalent increase in the fixation of beneficial AT alleles (Fig.2D and Fig. S2).

**Figure 2.**
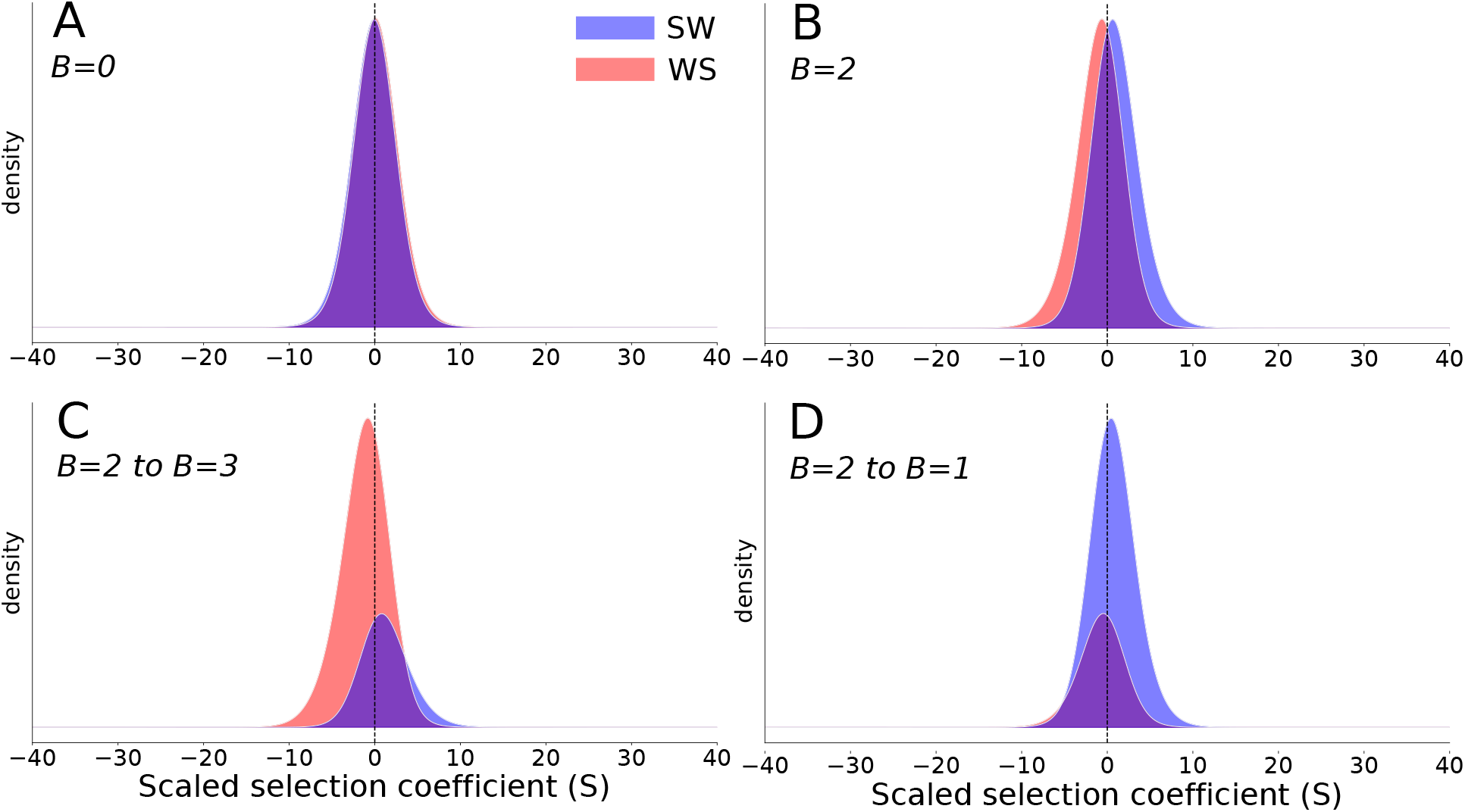
Distribution of fitness effects of new substitutions separately for WS (red) and SW (blue) substitutions under the concatenated DMS fitness landscape. A) Equilibrium frequencies were computed with *B* = 0 and the substitution rate also with *B* = 0 (no gBGC). B) Equilibrium frequencies were computed with *B* = 2 and the substitution rate also with *B* = 2. C) Equilibrium frequencies were computed with *B* = 2 and the substitution rate with *B* = 3 (increase in gBGC). D) Equilibrium frequencies were computed with *B* = 2 and the substitution rate with *B* = 1 (decrease in gBGC). The panel A&B therefore represent substitution rates at equilibrium, while panels C&D represent substitution rates out of equilibrium.

I then computed equilibrium codon frequencies under a wide range of gBGC intensity, mimicking genes evolving under different recombination rates. I then computed the DFE of substitutions in two scenarios. The first one corresponds to an increase of gBGC by 30% for all genes, the second to a decrease by the same amount. These scenarios mimic either a change in the genome-wide recombination rate, or a change in the meiotic repair bias. To visualize the displacement of the sequence in the fitness landscape, I computed the relative fitness of the sequence for all equilibrium states (see details in methods). I represented the proportions of beneficial, deleterious, WS, and SW substitutions in (Fig.3 row 1 and Fig. S3). When gBGC increases, genes that evolved under stronger gBGC show the higher proportion of deleterious substitutions towards GC (Fig.3C1 and Fig. S3). Conversely, when gBGC decreases, genes that previously evolved under gBGC show higher levels of beneficial substitutions towards AT than others (Fig.3A1 and Fig. S3).

**Figure 3.**
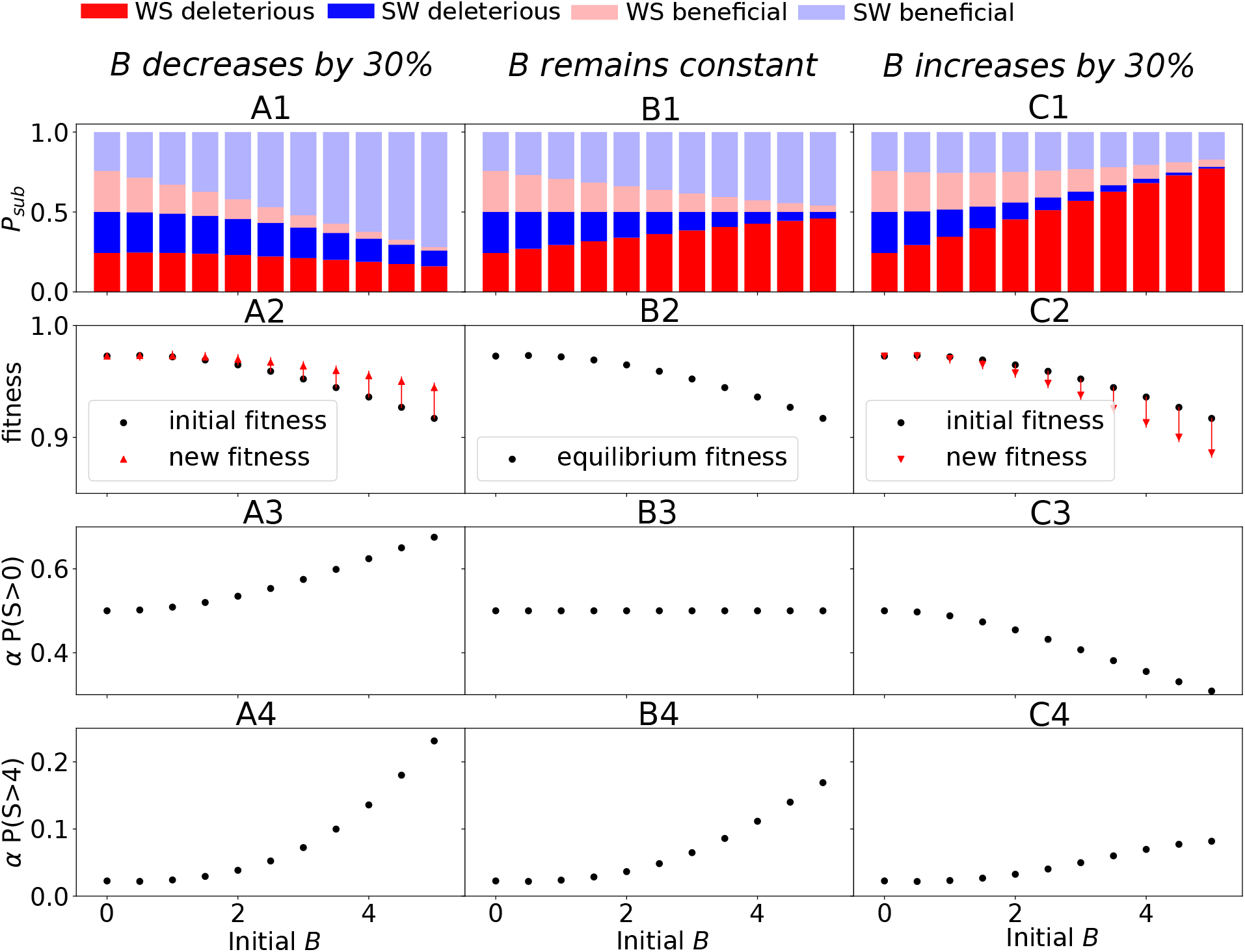
Row 1: Proportion of the substitutions (*P*_*sub*_) contributed by WS deleterious (bright red), SW deleterious (bright blue), WS beneficial (light red) and SW beneficial (light blue) mutations as a function of the population-scaled gBGC coefficient under the concatenated DMS fitness landscape in three scenarios: Equilibrium frequencies are computed with *B*, and substitutions from this equilibrium sequence are computed with 0.7 *× B* (A), *B* (B) and 1.3 *× B* (C). Row 2: Relative fitness (*f*) of sequences as a function of the *B* they are evolving under (black dots). Red arrows correspond to the fitness evolution of the sequences after A) a decrease of *B* by 30%, or C) after an increase of *B* by 30%. Row 3: Proportion of positively selected substitutions *P* (*S >* 0) in the three scenarios described above. Row 4: Proportion of strongly beneficial substitutions *P* (*S >* 4) in the three scenarios described above.

In population genetics, positive selection can be quantified by the parameter *α*, introduced by McDonald and Kreitman (1991), which is an estimator the proportion of positively selected substitutions (for all mutation types) among all substitutions. Here, because the fitness landscape is known, the proportion of positively selected substitutions is directly accessible from the DFE of substitutions. We can therefore investigate the effect of gBGC on the true proportion of positively selected substitutions that McDonald-Kreitman based methods aim to estimate. At equilibrium, without adaptation, we can see that *α* = 0.5. This is indeed expected at equilibrium, where the fitness is static. However, when gBGC increases, *α* decreases, substitutions are more deleterious, and the sequence falls down the fitness landscape (Fig.3C2&3 and Fig. S4). Conversely, when gBGC decreases *α* increases. This is due to the fact that a sequence that has previously experienced higher levels of gBGC climbs larger distances in the fitness landscape when gBGC is relaxed (Fig.3A2&3 and Fig. S4).

Usually, mutations with |*N*_*e*_*s*| *<* 1 are not included in *α*, since they are considered effectively neutral (Eyre-Walker, 2002). This is because *α* is usually intended to capture the proportion of adaptive substitutions, assuming that only adaptation (i.e. a change in the fitness landscape) can lead to strongly beneficial mutations (McDonald and Kreitman, 1991). Indeed, on a fixed fitness landscape and without gBGC, when effectively neutral mutations are discarded, *α ≃* 0 (Figure 3 row 4). However, at equilibrium, *α* increases with the strength of gBGC because gBGC induces the fixation of substitutions that are increasingly deleterious, and therefore compensated by substitutions that become strongly beneficial (Figure 3B4). When gBGC decreases, the increase of *α* with gBGC is higher than at equilibrium. This is because beneficial substitutions compensate the fixation of strongly deleterious mutations previously fixed under higher levels of gBGC while current levels of gBGC do not fix as many deleterious mutations (Figure 3A4). Finally, when gBGC increases, we still observe an increase of *α* with the strength of gBGC (lower than at equilibrium though) (Figure 3C4). While gBGC fixes more deleterious mutations than at equilibrium, the proportion of compensating strongly beneficial substitutions is still higher than without gBGC, where almost every substitution is effectively neutral. Therefore, paradoxically, genes that experience the highest *α* show the steeper decrease in fitness at the same time (Figure 3C2). The results presented above represent the expectation of the true impact of gBGC on positive selection and coding sequence evolution.

### Impact of gBGC fluctuations on the rate of coding sequence evolution

It has also been shown that gBGC can alter our ability to detect selection in protein-coding genes, mainly through incorrect interpretation of the *dN/dS* ratio (Berglund et al., 2009; Galtier et al., 2009; Ratnakumar et al., 2010; Bolívar et al., 2016). Originally, the *dN/dS* ratio was designed to quantify the selective constraints exerted on the sequence of a protein (Miyata and Yasunaga, 1980; Nei and Gojobori, 1986). It represents the probability of fixation of a mutation that alters the sequence of a protein, relative to a mutation that does not alter it, supposedly neutral (Spielman and Wilke, 2015). There are two key assumptions under which the *dN/dS* ratio serves its purpose: 1) allelic frequency changes of synonymous mutations are supposed to be due to drift only and 2) mutation rates of synonymous and non-synonymous changes within a gene are supposed to be identical, such that the ratio of substitution rates represents only differences in fixation probability (Spielman and Wilke, 2015). In the presence of gBGC, both those assumptions are violated: 1) synonymous mutations do not evolve only by drift but under a directional force that affects their substitution rate (Nagylaki, 1983), and 2) as the base composition differs at the three nucleotides of a codon (typically with the third codon showing higher GC-content in highly recombining genes), and SW mutation rates being usually higher that WS, mutation rates at synonymous and non-synonymous sites within a gene are different (Bolívar et al., 2016; Latrille and Lartillot, 2022). It appears therefore natural that the intensity of gBGC influences the *dN/dS* ratio (Berglund et al., 2009; Galtier et al., 2009; Ratnakumar et al., 2010; Bolívar et al., 2016).

First, I computed the *dN, dS* and *dN/dS* for sequences that reached equilibrium under *B* = 0 and that subsequently accumulate substitutions under *B* = 0 ↦ 10, mimicking the sudden birth of a recombination hotspot (Fig.4 and Fig. S5). Similarly, I computed *dN, dS* and *dN/dS* for sequences that reached equilibrium under *B* = 0 ↦ 10 and that subsequently accumulate substitutions under *B* = 0, mimicking the sudden death of a recombination hotspot (Fig.4 and Fig. S5).

**Figure 4.**
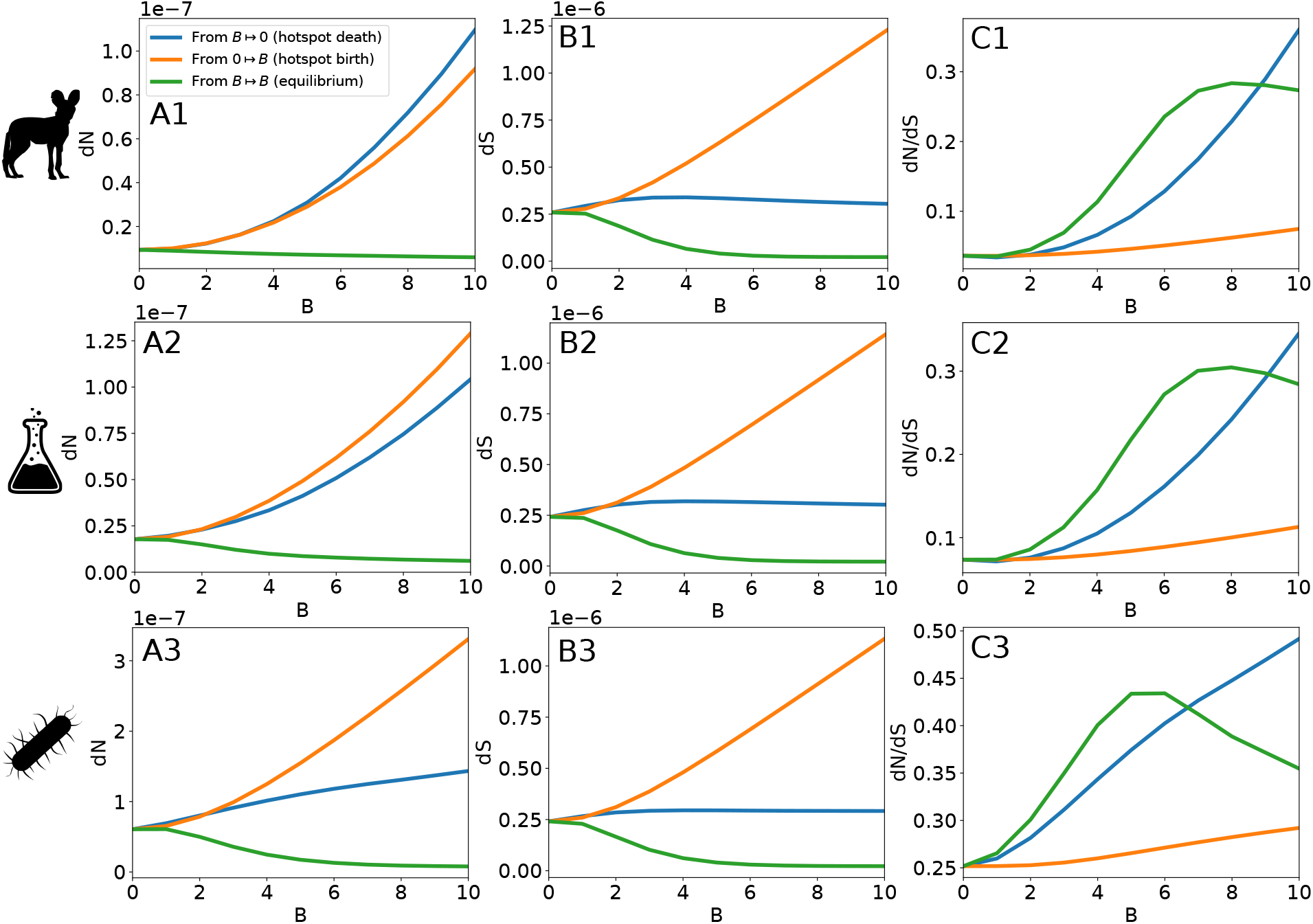
*dN* (A), *dS* (B), and *dN/dS* (C) as a function of the population scaled gBGC coefficient *B* in three scenarios: Equilibrium codon frequencies are computed without gBGC, and substitutions subsequently accumulate under a population-scaled gBGC coefficient of *B* (orange line), mimicking the birth of a recombination hotspot. Equilibrium codon frequencies are computed under a population-scaled gBGC coefficient of *B*, and substitutions subsequently accumulate without gBGC (blue line), mimicking the death of a recombination hotspot. And finally, equilibrium codon frequencies are computed under a population-scaled gBGC coefficient of *B*, and substitutions subsequently accumulate at equilibrium, under *B* (green line). From top to bottom: 2000 sites randomly sampled from the mammalian fitness landscapes, the concatenate of the DMS fitness landscapes (1389 sites) and the *E*.*coli* protein *β*-lactamase (263 sites).

The ratio *dN/dS* is higher at equilibrium than during the death of a hotspot (from *B* = 10 ↦ *B* = 0), than during its birth (from *B* = 0 ↦ *B* = 10) (Fig.4C). In both cases *dN* is high, either because of the fixation of deleterious GC alleles or because of the fixation of beneficial back-mutations towards AT (Fig.4A). The difference of *dN/dS* mainly stems from the different behaviour of the *dS* (Fig.4B). During the birth of a hotspot, *dS* is very high because of the rapid fixation of neutral GC alleles under strong gBGC. But when it dies, the increase of *dS* is small, and only due to the higher mutation rates of GC nucleotides, which are more abundant after a strong gBGC episode. Because the behaviour of the *dN/dS* is mainly the result of the impact of gBGC on the *dS*, the results presented here do not differ much with those of Bolívar et al. (2016) where the *dS* is modelled in the same way. However, Bolívar et al. (2016) predict a decrease of the *dN* when a sequence goes from a high GC content to a lower one (mimicking a decrease in gBGC), while the present model predicts an increase because of beneficial back-mutations. Overall, under all fitness landscapes, beneficial back-mutations affect the *dN/dS* ratio, and the death of a recombination hotspot induces a higher *dN/dS* than its birth.

### Non-synonymous GC-conservative substitution patterns are affected by gBGC

To avoid the confounding effect of gBGC on the *dN/dS* ratio, several studies computed *dN/dS* and *πN/πS* using GC-conservative substitutions/polymorphisms only (*G ↔ C* and *A ↔ T*) (Bolívar et al., 2016; Corcoran et al., 2017; Rousselle et al., 2019; Castellano et al., 2020). The idea being that as gBGC only distorts the segregation ratio of AT:GC heterozygous polymorphisms, the GC-conservative *dN/dS* should be an gBGC-robust estimator of the efficacy of purifying selection. However, although gBGC does not affect the fixation probability of GC-conservative mutations, by influencing the equilibrium sequence, it might affect mutational opportunities and therefore patterns of GC-conservative substitutions.

To explore this possibility, I also computed the DFE of WW and SS mutations with or without gBGC under all fitness landscapes. It appears that WW mutations are on average more deleterious in the presence of gBGC (*B*=2) under all fitness landscapes (Table S1). This is due to the fact that under gBGC, weakly constrained sites that are AT optimal are fixed in a GC state. Therefore, the only remaining opportunities for WW mutations are from sites that are sufficiently constrained to have resisted gBGC, and therefore these WW mutations are more deleterious on average. The consequence of this is a decrease of the WW *dN/dS* with recombination (Fig.5C and Fig. S6C). Importantly, since gBGC does not affect the fixation probability of WW mutations, this decrease depends only on past gBGC activity and not on the current value of gBGC. On the other hand, the impact of gBGC on the DFE of SS mutations is weak and depends on the fitness landscape (Table S1). In AT-optimal sites that have been fixed in a GC state because of gBGC, the DFE of SS mutations will depend on the fitness difference between the least deleterious GC-rich codon (favoured by selection) and the other deleterious GC-rich codons, which is not easily predictable and might differ depending on the fitness landscape. As a consequence the relationship between the SS *dN/dS* and recombination is weaker and sometimes non-monotonous (Fig.5C and Fig. S6C). As for WW substitutions, SS substitution patterns only depend on past gBGC activity. Overall, under realistic fitness landscapes, a slight decrease of the GC-conservative *dN/dS* ratio can in principle be caused by gBGC only (especially for WW mutations). Therefore, a negative correlation between recombination rate and the GC-conservative *dN/dS* does not necessarily reflect an increased efficacy of purifying selection as proposed by Bolívar et al. (2016), Corcoran et al. (2017), Rousselle et al. (2019) and Castellano et al. (2020). Importantly, under the assumptions of the model, although the rates of non-synonymous WW and SS substitutions are affected by gBGC, the proportions of positively selected WW and SS substitutions remain fairly unaffected (Fig. S7).

**Figure 5.**
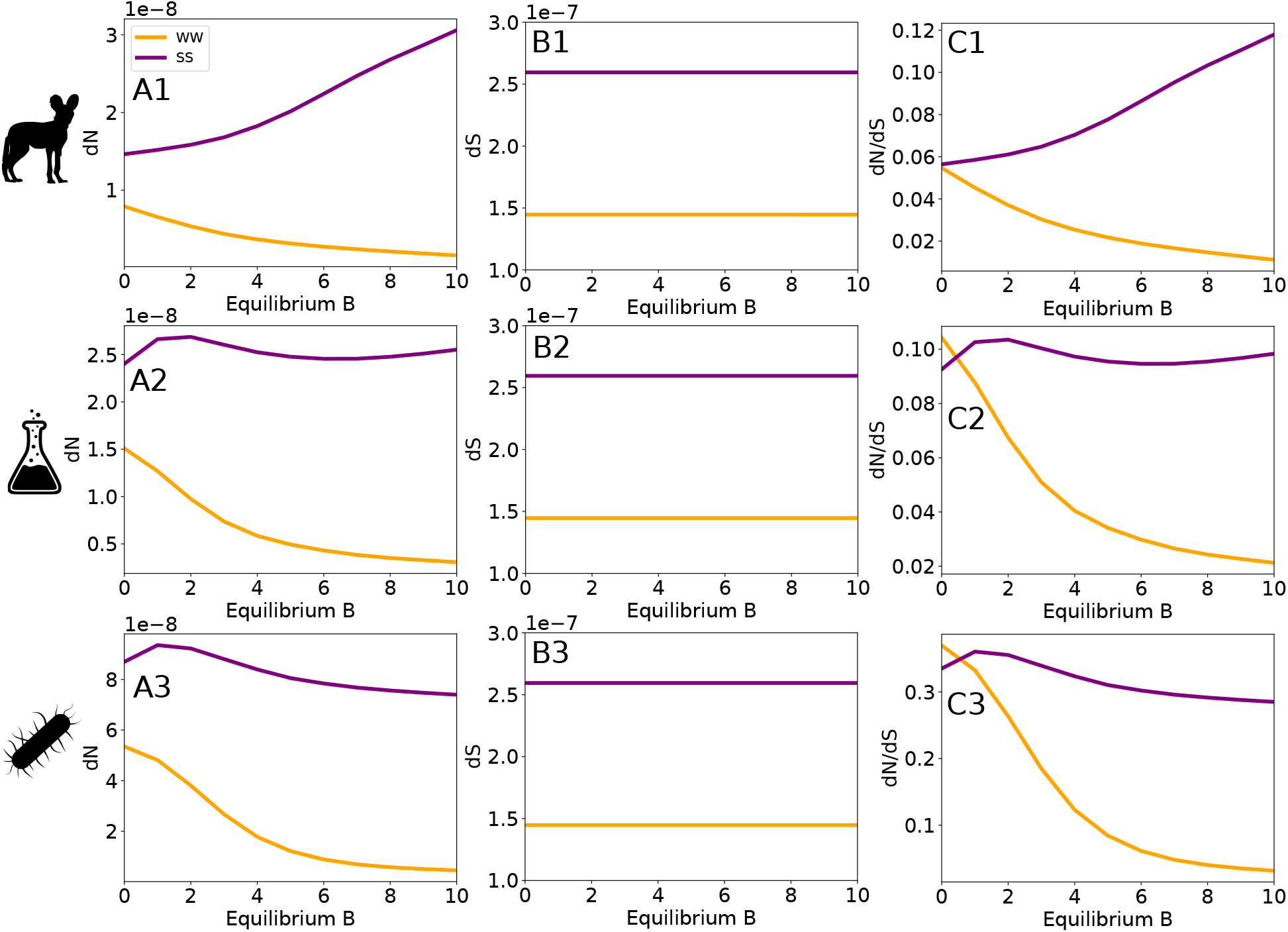
*dN* (A), *dS* (B), and *dN/dS* (C) as a function of the equilibrium population-scaled gBGC coefficient *B* for WW and SS substitutions. From top to bottom: 2000 sites randomly sampled from the mammalian fitness landscapes, the concatenate of the DMS fitness landscapes (1389 sites) and the *E*.*coli* protein *β*-lactamase (263 sites).

## Discussion

In this study, I estimated the impact of beneficial back-mutations on the dynamics of natural selection in the presence of gBGC. I showed that gBGC induces a shift towards positive values in the DFE of new mutations. When the intensity of gBGC decreases, this leads to an increase of beneficial substitutions, and a decrease of deleterious ones. If not properly investigated, this selection dynamic can be misinterpreted as a beneficial effect of recombination, when paradoxically it is fuelled by its deleterious consequences. Moreover, I showed that one can also expect an increase of the *dN/dS* due to these beneficial-back mutations. These results strengthen those of previous studies demonstrating that in the presence of gBGC, the *dN/dS* ratio is not a correct estimator of the selective pressure exerted on proteins (Berglund et al., 2009; Galtier et al., 2009; Ratnakumar et al., 2010; Bolívar et al., 2016, 2019; Rousselle et al., 2019). Of note, I show that the GC-conservative *dN/dS*, previously thought to be a gBGC-robust estimator of the efficacy of natural selection, is in fact also affected by gBGC.

Moreover, the present result highlights that the absence of a substitution pattern skewed towards GC, affecting both coding and non-coding sequences does not allow the interpretation of local accelerated evolution as an adaptation to changing environments. It appears therefore unwise to infer adaptive evolution from the *dN/dS* ratio of a given gene without prior information on gBGC, the fitness landscape, the local recombination dynamic or an external corroboration (e.g. correlation with phenotypic evolution or environmental changes).

This being said, it is still very possible that Hill-Robertson interference dissipation in highly recombining genes helps to fix beneficial mutations. In this sense, some studies have reported a positive correlation between measures of positive selection and recombination in *Drosophila melanogaster* (Castellano et al., 2016; Murga-Moreno et al., 2019), where the contrast between lowly recombining genes and highly recombining ones is higher than in humans (Bullaughey et al., 2008), and where the current levels of gBGC seem to be very low (Robinson et al. (2014) but see Jackson and Charlesworth (2021)).

### Comparison with empirical results

In several studies, it was observed that genes richer in GC3 (supposedly having experienced higher recombination rates) have a lower *dN/dS*. Some of these studies interpreted this pattern as higher selection efficacy due to the dissipation of Hill-Robertson interference (Gossmann et al., 2014; Rousselle et al., 2019), while another argued that it was a consequence of gBGC (Bolívar et al., 2016). The rationale behind the latter claim is that when a sequence is strongly constrained, the *dN* will be little affected by gBGC while the *dS* will rise quickly, leading to a lower *dN/dS* (Bolívar et al., 2016). Here, we do not observe a regime in which the *dN/dS* decreases, because the *dS* decreases always faster than the *dN*, leading to an increase of the *dN/dS*. However, both the present model and that of Bolívar et al. (2016) predict a decrease of the *dS* at equilibrium, while empirical data suggest an increase (Bolívar et al., 2016; Corcoran et al., 2017; Rousselle et al., 2019). This increase is usually attributed to a mutagenic effect of recombination (Bolívar et al., 2016; Rousselle et al., 2019; Castellano et al., 2020), not modelled here, although there are many properties that are correlated with GC3 that can explain an increase of the synonymous divergence. It is indeed difficult to interpret correlations between *dN/dS* and GC3 in empirical data, because genes with different GC3 might also evolve under different fitness landscapes (they tend to have different functions, see Pouyet et al. (2017)) and different mutational processes (they might be more susceptible to CpG hyper-mutability) that are not directly linked to gBGC or recombination.

### The equilibrium assumption in regards of hotspot dynamics

In this study, for mathematical convenience, I chose to model out of equilibrium dynamics by first computing codon frequencies under equilibrium conditions and then computing substitution rates under new conditions. In mammals, for instance, as most of the genome has a rather low recombination rate, it is reasonable to say that most genes are around equilibrium for a rather low strength of gBGC. In this case, the birth of a strong hotspot will indeed leave a clear signature of gBGC in substitution patterns (Berglund et al., 2009; Galtier et al., 2009; Ratnakumar et al., 2010; Clément and Arndt, 2013; Lesecque et al., 2014). However, in humans and mice, the vast majority of hotspots are short-lived (Auton et al., 2012; Smagulova et al.,2016; Pratto et al., 2014; Alleva et al., 2021). Therefore, it is very likely that these short-lived hotspots do not have time to reach equilibrium, and the signatures of the death of a hotspot in the *dN/dS* may not be strong enough to be observed in individual hotspots in these species. Of note, those hotspot still remain long enough so that they leave a clear gBGC footprint in substitutions across mammals (Berglund et al., 2009; Galtier et al., 2009; Ratnakumar et al., 2010; Clément and Arndt, 2013; Lartillot, 2013; Lesecque et al., 2014; Galtier, 2021). On the other hand, many species including most placental mammals, passerine birds, colubroid snakes, budding yeasts and many angiosperm plants exhibit recombination hotspots in 5’ of genes that are relatively long-lived (Axelsson et al., 2012; Choi and Henderson, 2015; Singhal et al., 2015; Lam and Keeney, 2015; Kawakami et al., 2017; Schield et al., 2020; Hoge et al., 2023; Joseph et al., 2023). Interestingly, in mammals these recombination hotspots co-evolve slowly with DNA methylation (Joseph et al., 2023). When those long-lived hotspots stop to be hypomethylated, they usually die (Joseph et al., 2023). Thus, they should leave a strong signal of positive selection, that do not correspond to an adaptive response of the gene to changing environments.

Altogether, both fast and slow hotspot dynamics should lead to pervasive fixation of deleterious mutations because of gBGC, compensated by positive selection. Even if this positive selection can be weak at a given gene when hotspots are short-lived, it should still have a significant impact throughout the genome.

### Beneficial back-mutations versus compensatory mutations

The fitness landscapes used in this study are simplistic, in the sense that they imply that each aminoacid evolves independently, and does not interact with others (no epistasis). However, numerous studies found that epistasis plays an important role in protein evolution (Bonhoeffer et al., 2004; Breen et al., 2012; Starr and Thornton, 2016; Miton and Tokuriki, 2016). Let us consider the simple example of two residues interacting such that the optimal state of the protein requires either an alanine at one site and a valine at the other, or a lysine at one site and an arginine at the other. If the first residue mutates from alanine to lysine, optimal protein state can either be restored by a beneficial back-mutation from lysine to alanine, or a compensatory mutation at the other site from valine to arginine. In this simple case, it is easy to see that the fixation of a deleterious amino-acid does not always lead to an opportunity for positive selection at the same site. In the end, with epistasis, the deleterious effect of gBGC at one site/gene can be compensated by positive selection at another site/gene. Of note, if compensatory mutations occur preferentially in the same gene, one still expects a decrease of the genome-wide strength of gBGC to increase positive selection in highly recombining genes.

Interestingly, deep mutational scanning experiments showed that in some proteins, the site-specific fitness landscape can remain unchanged even in divergent lineages and predicts quite accurately the evolution of sequences *in natura* (Ashenberg et al., 2013; Doud et al., 2015; Bloom, 2017). This suggests that contrarily to the previous example, fitness at one site does not always depends on the amino-acids present at other sites. In this sense, there is also accumulating evidence for convergent adaptation at the molecular level (Christin et al., 2007; Zhen et al., 2012; Davies et al., 2012; Wu et al., 2020; Duchemin et al., 2023; Fukushima and Pollock, 2023), suggesting that there is sometimes a limited number of mutations that can lead to a given phenotype, and thus limited opportunities for compensatory mutations. Moreover, an important role of beneficial back mutations have been reported in plants (Chen et al., 2021) and mammals (Moses and Durbin, 2009; Latrille et al., 2023). A recent study suggests that beneficial-back mutations constitute an important fraction of beneficial mutations in several mammals (between 20% and 40%) (Latrille et al., 2023). Finally, several human accelerated genes show a remarkably strong AT-biased substitution pattern (Galtier et al., 2009), but they have not been more deeply investigated, and have been left unchecked in other studies (Ratnakumar et al., 2010; Kostka et al., 2012).

Altogether, while epistasis can spread throughout the genome the compensatory response to a strong gBGC episode, this compensation might not always be possible, and beneficial back-mutations are therefore expected to leave a signature of positive selection upon a decrease of gBGC. Still, the relative contribution of beneficial back-mutations and compensatory mutations to stabilizing selection in general is an open question that remains to be investigated, and the burst of fixation of deleterious mutations induced by gBGC could in fact be an interesting case study to investigate it.

### Incorporating beneficial back-mutations into evolutionary thinking

In population genetics, the infinite site model approximation inherently excludes the possibility of beneficial back-mutations (Kimura, 1969; Ohta, 1992). This approximation can be very reasonable mathematically at a very short timescale, and is widely used in population genetics. Nevertheless, all sequences are the product of a very long evolutionary history. With a finite genome and longer evolutionary time, beneficial back-mutations are expected to occur (Gillespie, 1995; Hartl and Taubes, 1996; Piganeau and Eyre-Walker, 2003; Sella and Hirsh, 2005; Charlesworth and Eyre-Walker, 2007; Mustonen and Lässig, 2009). Under a slowly evolving fitness landscape, they largely contribute to molecular evolution (Chen et al., 2021; Latrille et al., 2023). Modelling evolution as the displacement of a sequence on a fitness landscape, determined by natural selection and nonadaptive evolutionary forces, instead of modelling selection with a constant DFE is therefore more relevant to study selection dynamics (Halpern and Bruno, 1998; Rodrigue et al., 2010; Tamuri et al., 2012; Rodrigue and Lartillot, 2014; Jones et al., 2017; Tamuri and dos Reis, 2022; Latrille et al., 2023).

Using this framework, I showed that signatures of positive and negative selection cannot be used to conclude on a beneficial or negative effect of recombination. For species where meiotic gene conversion is biased towards GC, this seriously questions all the previous interpretations of this beneficial effect (or its absence) that are based on the *dN/dS* or other methods contrasting non-synonymous and synonymous changes (Bullaughey et al., 2008; Gossmann et al., 2014; Castellano et al., 2016; Bolívar et al., 2016; Rousselle et al., 2019; Murga-Moreno et al., 2019; Grandaubert et al., 2019; Hämälä and Tiffin, 2020; Castellano et al., 2020; Cavassim et al., 2021). In presence of gBGC, it is thus important to account for beneficial back-mutations.

Even without gBGC, this work is consistent with previous theoretical studies showing that the nearly universal mutation bias towards AT necessarily induces a fixation bias towards GC in selectively constrained sequence, also because of beneficial back-mutations (Latrille and Lartillot, 2022; Kaj et al., 2023). This calls for a re-interpretation of results from several studies that interpreted the fact that GC alleles segregate at higher frequency at non-synonymous sites as an evidence for the negative impact of gBGC on adaptation (Hämälä and Tiffin, 2020; Liang et al., 2022). For the same reasons that increased positive selection in highly recombining genes does not necessarily reflect a beneficial effect of recombination, a fixation bias towards GC at non-synonymous sites does not necessarily reflect the presence of gBGC.

Incorporating beneficial back-mutations into evolutionary thinking is therefore essential for understanding selection dynamics in the presence of biases, and should prevent the spread of many misinterpretations and erroneous conclusions.

## Material and methods

### Fitness landscapes

The mammalian fitness landscape was reconstructed by fitting a mutation-selection model to a multispecies alignment of 14,509 protein-coding genes of 87 mammalian species in Latrille et al. (2023). This fitness landscape is site-specific: the fitness of amino-acids at one site does not depend on amino-acids at other sites (epistasis is neglected). Importantly, mutation-selection models cannot disentangle selection coefficients from effective population sizes (Rodrigue et al., 2010). The model can only estimate scaled fitness differences, assuming a constant effective population size throughout the mammalian evolutionary history. The experimental fitness landscapes were taken from Deep mutational scanning studies on the virus Influenza (Thyagarajan and Bloom, 2014; Doud et al., 2015), on the bacteria *E*.*coli* (Stiffler et al., 2015) and on the yeast *S*.*cervisae* (Kitzman et al., 2015). Except for yeasts, the experimental fitness landscapes have therefore not been obtained in organisms that are particularly subject to gBGC (Mancera et al., 2008), but still reflect selective pressures exerted on proteins in general. These experimental fitness landscapes were retrieved from the study of Bloom (2017). Individual proteins with peculiar functions can have peculiar fitness landscapes which are not necessarily representative of the genome. To have a beginning of an average fitness landscape, I also performed analyses on a concatenate of the four DMS fitness landscape.

### Codon equilibrium frequencies

Using a Wright-Fisher diffusion approximation, for a given site *l*, one can compute the substitution rates from codon *i* to codon 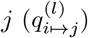 (Nagylaki, 1983; Halpern and Bruno, 1998).

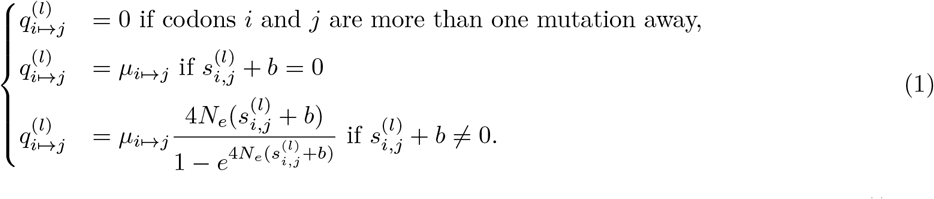

Here, *µ*_*i* ↦ *j*_ is the mutation rate from codon *i* to codon *j, N*_*e*_ the effective population size, 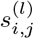 the difference of fitness between codon *i* and codon *j* at site *l*, and *b* the gBGC coefficient. When codon *i* and *j* are separated by a synonymous mutation, I consider that *s*_*i,j*_ = 0. The gBGC coefficient is the product of the repair bias towards GC, the recombination rate per base pair and the length of the conversion tract.

A local increase of recombination rate thus directly increases the intensity of gBGC. If the mutation that separates codon *i* from codon *j* is from AT to GC (WS), *b* is positive, if it is from GC to AT (SW), *b* is negative, and if it is GC conservative (A *↔* T or C *↔* G), *b* = 0. Two mutation matrices have been used: a Jukes-Cantor mutation matrix where all mutation rates are equal, and an empirical mutation matrix estimated from singletons of the human’s chromosome 1 extracted from the human 1000 genomes project (Gazal et al., 2015).

One can then build a 61x61 transition matrix for each pair of codons ***Q***^**(*l*)**^. The diagonal of this matrix is the negative sum of the *q*_*i,j*_:

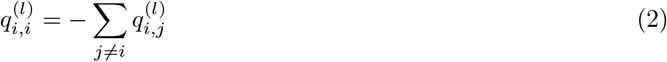

From this transition matrix, the vector of equilibrium frequencies of each codon ***X***^**(*l*)**^ can be calculated by solving the following system:

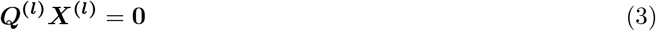

### DFE of new mutations

At equilibrium, the probability for observing a mutation from codon *i* to codon *j* is 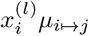 where 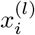 is the equilibrium frequency of codon *i* at site *l*. For the whole sequence, the DFE of new mutations represented in Fig.1 has been computed as:

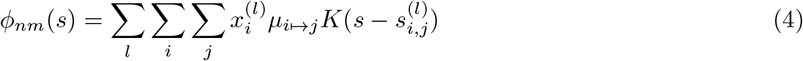

where *K* is a Gaussian kernel function.

### DFE of substitutions

Similarly to the DFE of new mutations, at equilibrium, the probability of observing a substitution from codon *i* to codon *j* is 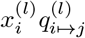. For the whole sequence, the DFE of substitutions represented in Fig.2 has been computed as:

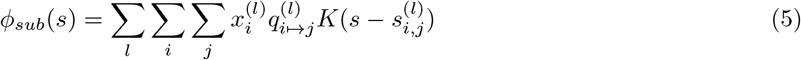

### Relative fitness

The relative fitness at a given site is defined as the average fitness of all codons weighted by their equilibrium frequency, divided by the fitness of the fittest codon. The relative fitness of the sequence *f* is just the average of the relative fitnesses of all sites:

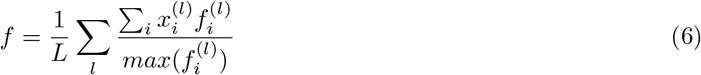

where 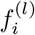 is the fitness of codon *i* at site *l*, and *L* the total number of sites.

### dN/dS

I computed *dN/dS* as the ratio between the rates of non-synonymous over synonymous substitutions given by:

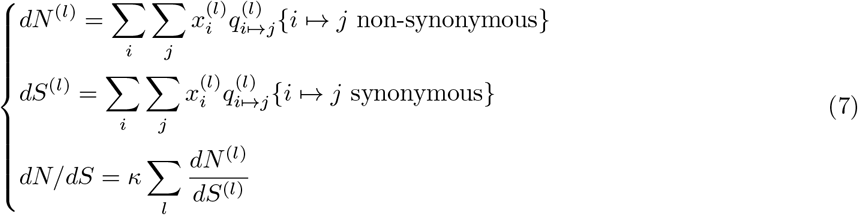

where *κ* is a constant normalisation factor corresponding to the ratio of the number of possible synonymous mutations over the number of possible non-synonymous mutations from the equilibrium sequence.

## Acknowledgments

I wish to thank Philippe Veber and Thibault Latrille for insightful discussions on mutation-selection models and Nicolas Lartillot, Carina Farah Mugal, Laurent Duret and Bastien Boussau for their very helpful reviews on a first version of this manuscript.

## Data and materials availability

Analysis scripts and documentation are available at https://gitlab.in2p3.fr/julien.joseph/backdfe

