## Supplementary material for "Increased positive selection in highly recombining genes does not necessarily reflect an evolutionary advantage of recombination"

### Supplementary material from: Increased positive selection in highly recombining genes does not necessarily reflect a beneficial effect of recombination

**J. Joseph**[1]

[1]LBBE, Université Lyon 1, CNRS, UMR 5558, Villeurbanne, France

April 8, 2024

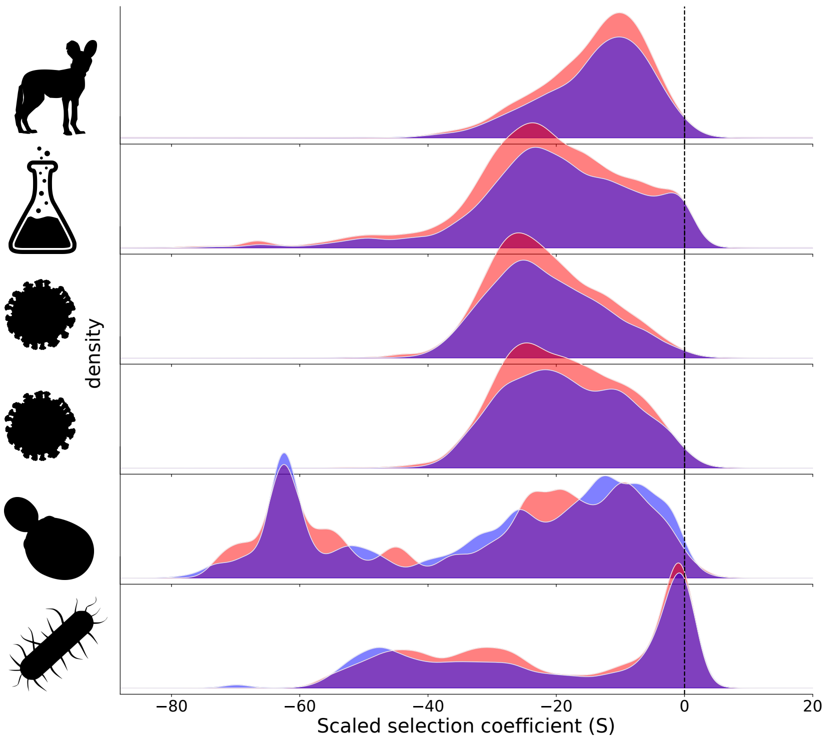


Figure S1: Distribution of fitness effects of new mutations at equilibrium separately for WS (red) and SW (blue) mutations. For each fitness landscape, the relative fitnesses were shuffled among amino-acids. Equilibrium frequencies were computed with $B = 0$. From top to bottom: 2000 sites randomly sampled from the mammalian fitness landscapes, the concatenate of the DMS fitness landscapes (1389 sites), the fitness landscape of the influenza protein NP (498 sites), the influenza protein HA (564 sites), the *S.cervisae* protein Gal4 (64 sites) and the *E.coli* protein $\beta$-lactamase (263 sites).

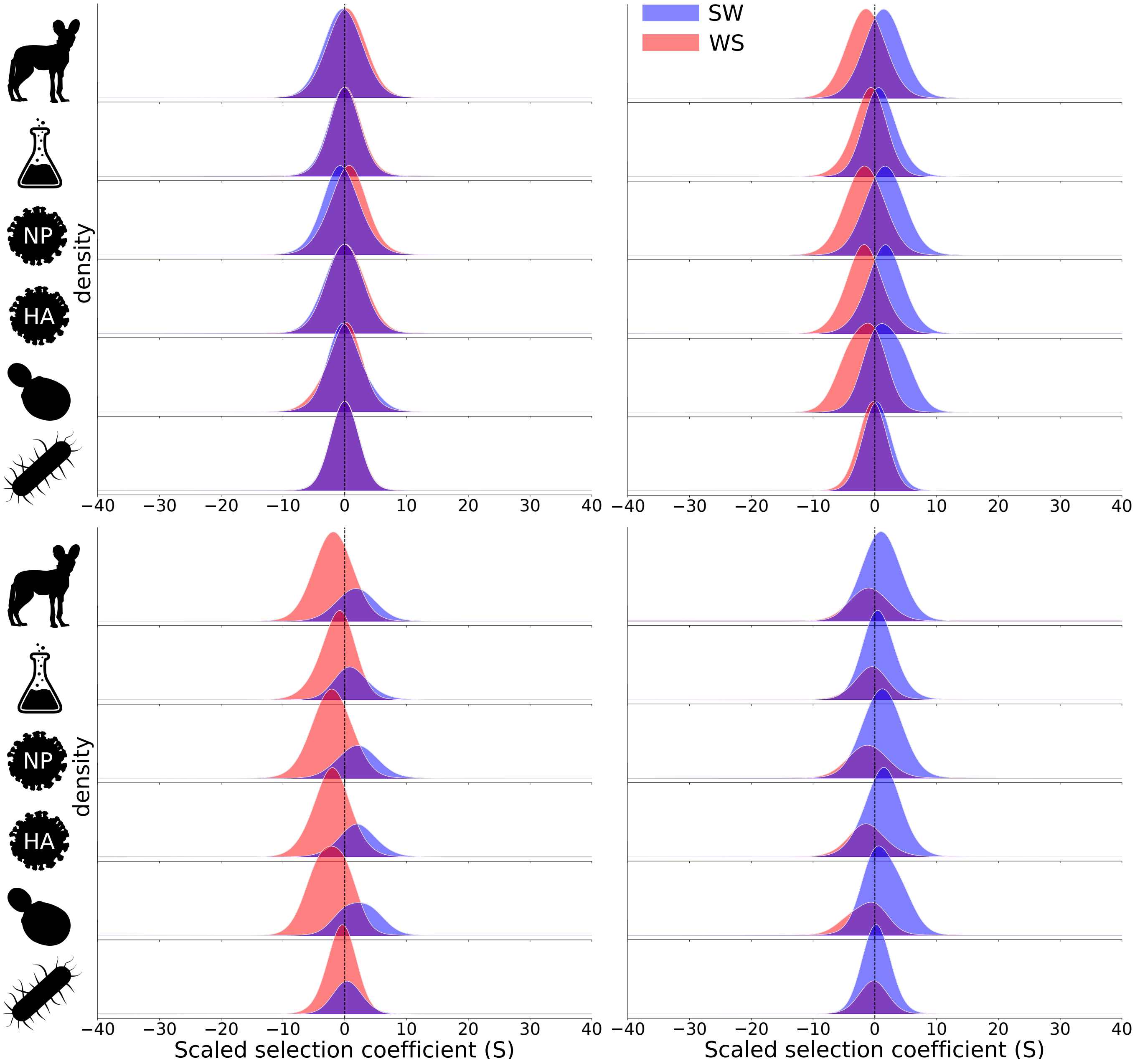


Figure S2: Distribution of fitness effects of new substitutions separately for WS (red) and SW (blue) substitutions. Equilibrium frequencies were computed for the concatenated DMS fitness landscape, with $B = 0$ (top left) and $B = 2$ (top right and bottom). The substitution rate from the equilibrium sequence was subsequently computed with $B = 0$ (top left), $B = 2$ (top right), $B = 3$, (bottom left), and $B = 1$ (bottom right). The top panels therefore represent substitution rates at equilibrium, while the bottom panels represent substitution rates out of equilibrium. From top to bottom: 2000 sites randomly sampled from the mammalian fitness landscapes, the concatenate of the DMS fitness landscapes (1389 sites), the fitness landscape of the influenza protein NP (498 sites), the influenza protein HA (564 sites), the *S.cervisae* protein Gal4 (64 sites) and the *E.coli* protein $\beta$-lactamase (263 sites).

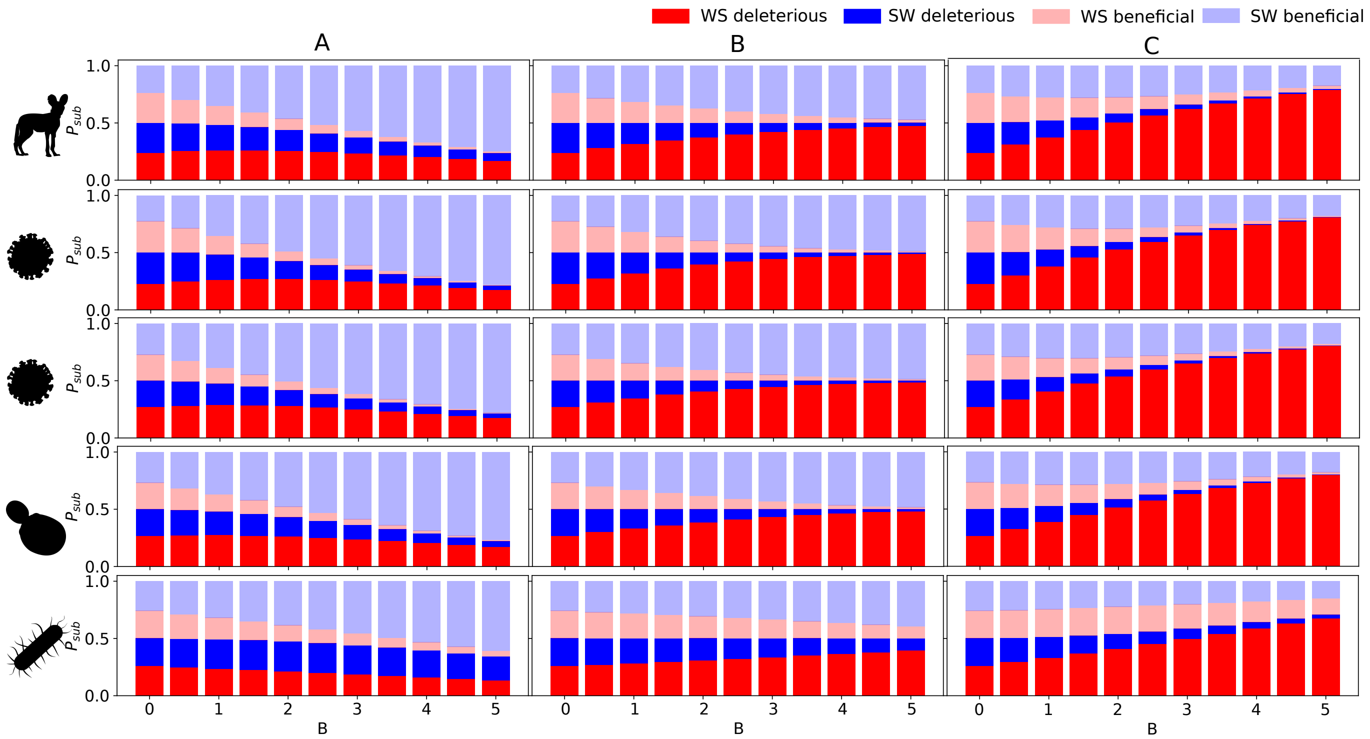


Figure S3: Proportion of the substitutions ($P_{sub}$) contributed by WS deleterious (bright red), SW deleterious (bright blue), WS beneficial (light red) ans SW beneficial (light blue) mutations as a function of the population-scaled gBGC coefficient under the concatenated DMS fitness landscape in three scenarios: Equilibrium frequencies are computed with $B$, and substitutions from this equilibrium sequence are computed with $0.7 \times B$ (A), $B$ (B) and $1.3 \times B$ (C). From top to bottom: 2000 sites randomly sampled from the mammalian fitness landscapes, the fitness landscape of the influenza protein NP (498 sites), the influenza protein HA (564 sites), the *S.cervisae* protein Gal4 (64 sites) and the *E.coli* protein $\beta$-lactamase (263 sites).

| | **B = 0 JC** | | **B = 0 HM** | | **B = 2 HM** | |
|---|---|---|---|---|---|---|
| **Fitness landscape** | **WW** | **SS** | **WW** | **SS** | **WW** | **SS** |
| **Mammalian** | 1.29% | 1.59% | 1.47% | 1.55% | 0.98% | 1.67% |
| **Concatenated DMS** | 2.99% | 3.27% | 3.32% | 2.98% | 2.09% | 3.4% |
| **NP (Influenza)** | 0.53% | 0.66% | 0.55% | 0.66% | 0.50% | 0.63% |
| **HA (Influenza)** | 1.42% | 1.47% | 1.56% | 1.46% | 1.18% | 1.41% |
| **Gal4 (*S.cervisae*)** | 1.96% | 1.23% | 2.24% | 1.03% | 1.35% | 1.36% |
| **$\beta$-lactamase (*E.coli*)** | 12.09% | 12.77% | 12.71% | 11.97% | 9.13% | 12.86% |

Table S1: Proportion of beneficial mutations under the two equilibrium conditions $B = 0$ and $B = 2$ for all the fitness landscapes of the study.

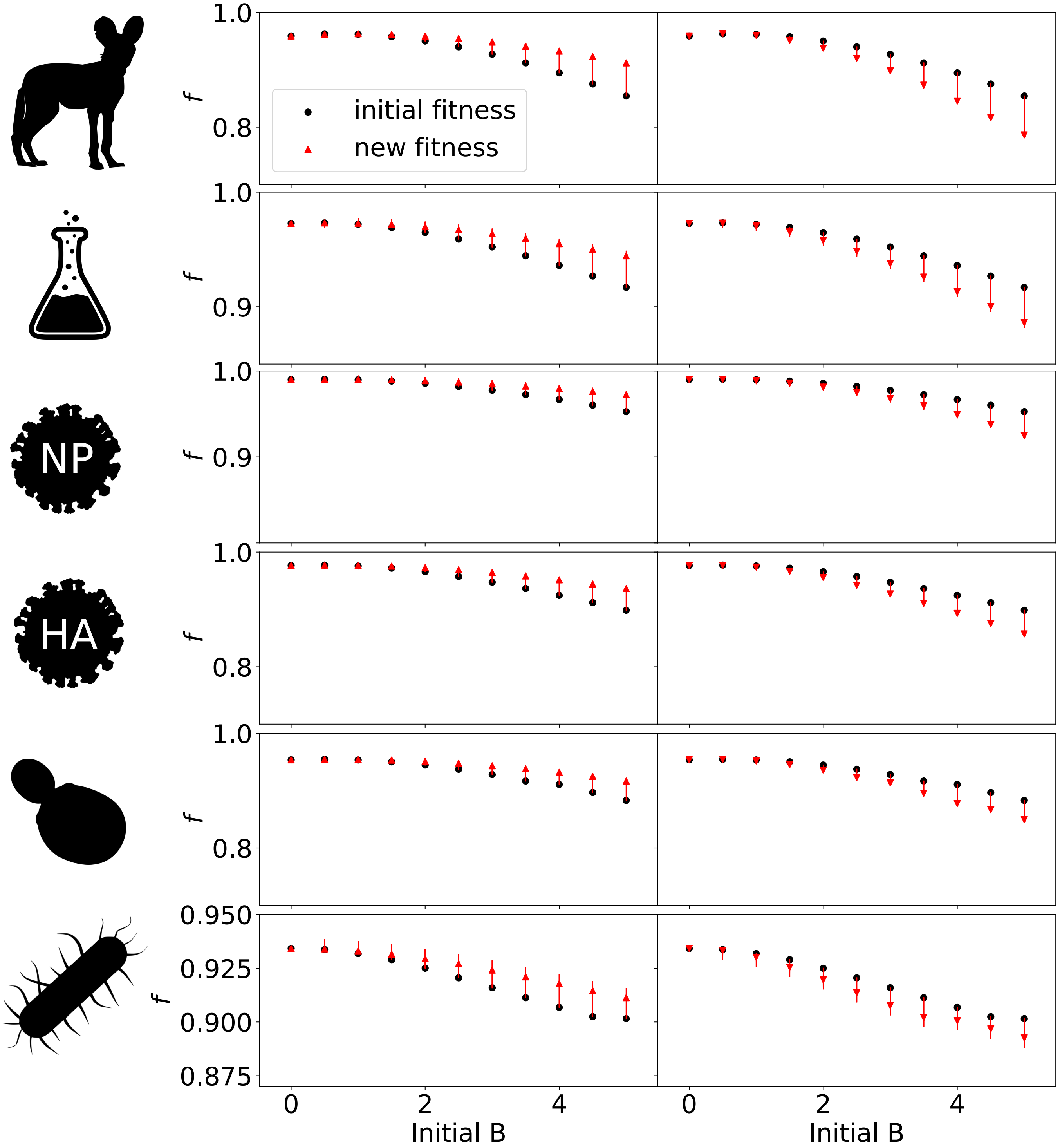


Figure S4: Relative fitness ($f$) of sequences as a function of the $B$ they are evolving under (black dots). Red arrows correspond to the fitness evolution of the sequences after a decrease of $B$ by 30% (left panels), or after an increase of $B$ by 30% (right panels). From top to bottom: 2000 sites randomly sampled from the mammalian fitness landscapes, the fitness landscape of the influenza protein NP (498 sites), the influenza protein HA (564 sites), the *S.cervisae* protein Gal4 (64 sites) and the *E.coli* protein $\beta$-lactamase (263 sites).

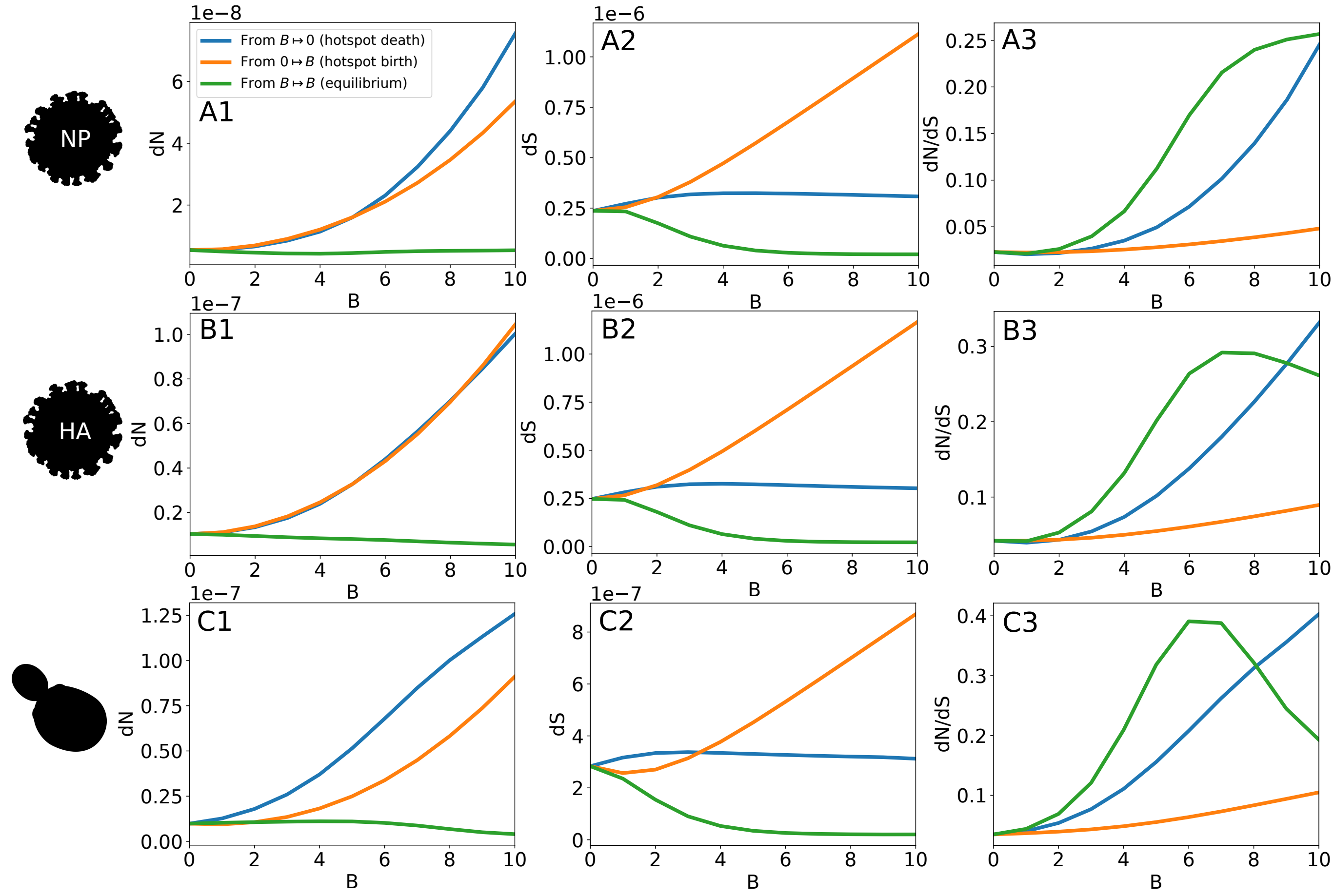


Figure S5: $dN$ (A), $dS$ (B), and $dN/dS$ (C) as a function of the population scaled gBGC coefficient $B$ in three scenarios: Equilibrium codon frequencies are computed without gBGC, and substitutions subsequently accumulate under a population-scaled gBGC coefficient of $B$ (orange line), mimicking the birth of a recombination hotspot. Equilibrium codon frequencies are computed under a population-scaled gBGC coefficient of $B$, and substitutions subsequently accumulate without gBGC (orange line), mimicking the death of a recombination hotspot. And finally, equilibrium codon frequencies are computed under a population-scaled gBGC coefficient of $B$, and substitutions subsequently accumulate at equilibrium, under $B$ (green line). From top to bottom: the fitness landscape of the influenza protein NP (498 sites), the influenza protein HA (564 sites) and the *S.cervisae* protein Gal4 (64 sites).

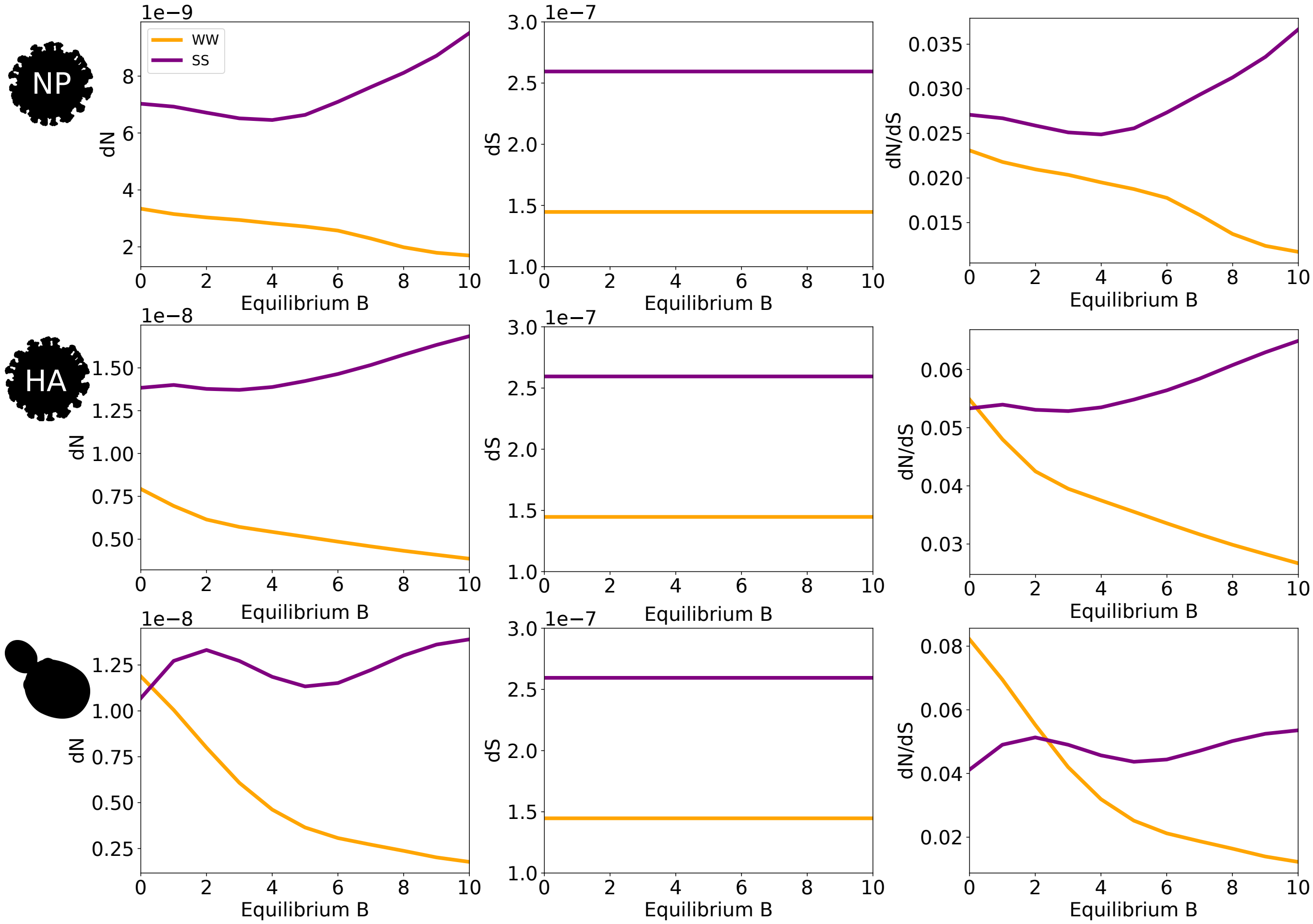


Figure S6: $dN$, $dS$, and $dN/dS$ as a function of the equilibrium population scaled gBGC coefficient $B$ for WW and SS substitutions. From top to bottom: the fitness landscape of the influenza protein NP (498 sites), the influenza protein HA (564 sites) and the *S.cervisae* protein Gal4 (64 sites).

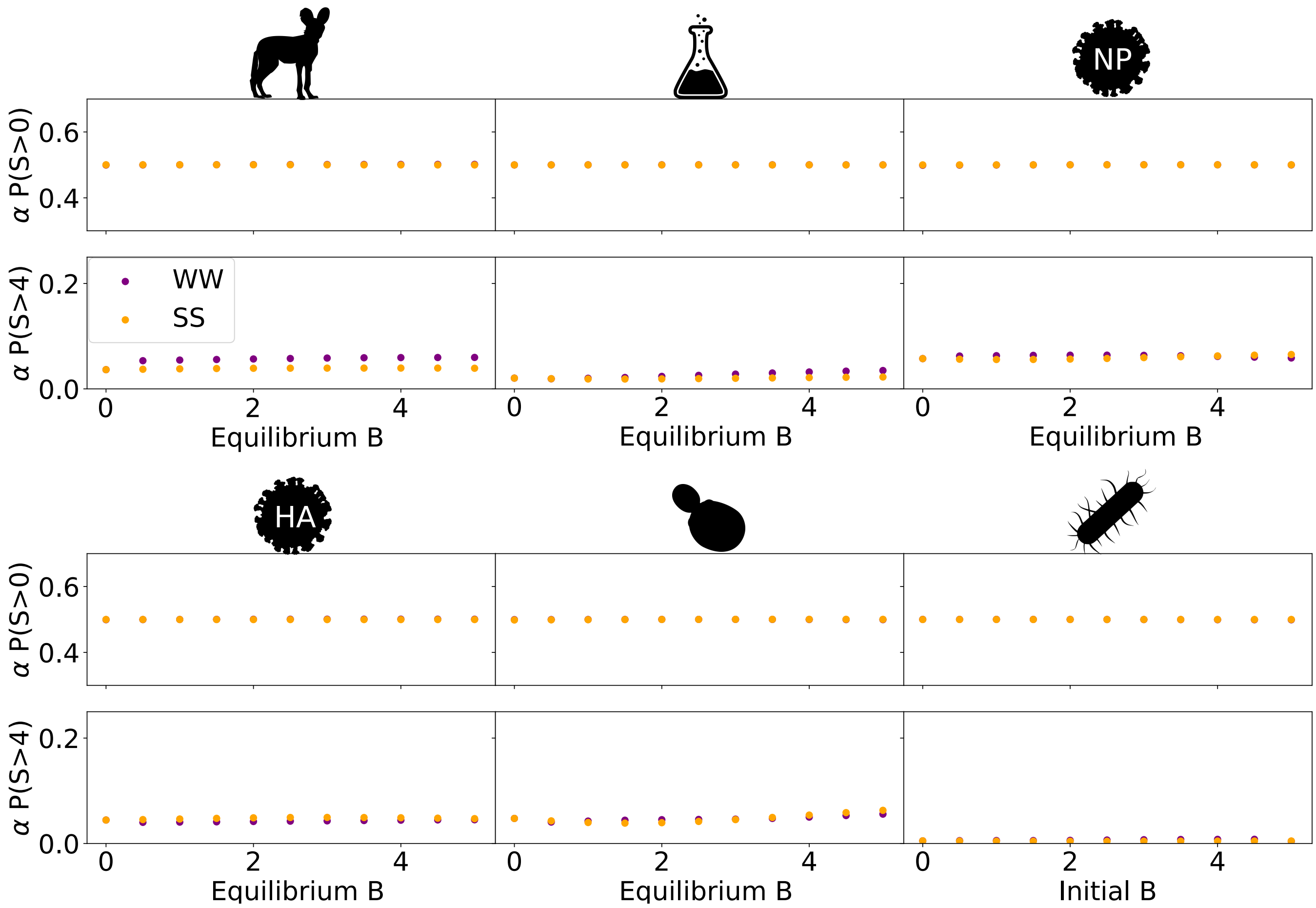


Figure S7: Proportion of positively selected WW (purple) and SS (orange) substitutions $P(S > 0)$ (first row) and $P(S > 4)$ (second row) as a function on the equilibrium population-scaled gBGC coefficient ($B$) for all fitness landscapes.
